# Overdominant and partially dominant mutations drive short-term adaptation in diploid yeast

**DOI:** 10.1101/2021.10.29.466440

**Authors:** Dimitra Aggeli, Daniel A. Marad, Xianan Liu, Sean W. Buskirk, Sasha F. Levy, Gregory I. Lang

## Abstract

Identification of adaptive targets in experimental evolution typically relies on extensive replication and allele reconstructions. An alternative approach is to directly assay all mutations in an evolved clone by generating pools of segregants that contain random combinations of the evolved mutations. Here, we apply this method to 6 clones isolated from 4 diploid populations that were clonally evolved for 2,000 generations in rich glucose medium. Each clone contains ∼20-25 mutations relative to the ancestor. We derived intermediate genotypes between the founder and the evolved clones by bulk mating sporulated cultures of each evolved clone to a barcoded haploid version of the founder. We competed the barcoded segregants *en masse* and quantified the fitness of each barcode. We estimated average fitness effects of evolved mutations using barcode fitness and whole genome sequencing for a subset of segregants or time-course whole population whole genome sequencing. In contrast to our previous work in haploid populations, we find that diploids carry fewer evolved mutations with a detectable fitness effect (6%), contributing a modest fitness advantage (up to 5.4%). In agreement with theoretical expectations, reconstruction experiments show that all adaptive mutations manifest some degree of dominance over the ancestral allele, and most are overdominant. Competition assays under conditions that deviated from the evolutionary environment show that adaptive mutations are often pleiotropic.

## INTRODUCTION

Over the course of adaptation of a clonal population, selection acts on genetic variation that is generated within the first few hundred generations (Lang, Botstein, and Desai 2011; Blundell et al. 2019). The fates of these mutations are not independent, and many beneficial mutations will be lost due to genetic drift and clonal interference (Lang, Botstein, and Desai 2011; Good et al. 2017). In addition, genetic hitchhiking and mutational cohorts will result in the fixation of many neutral or deleterious mutations (Lang et al. 2013). Collectively these effects make it difficult to determine which mutations in an evolved clone are adaptive (Lang, Botstein, and Desai 2011; Lang et al. 2013; Buskirk, Peace, and Lang 2017; Kvitek and Sherlock 2011). One possibility is to leverage replicate cultures in order to identify mutations in common targets (Lang et al. 2013; Marad, Buskirk, and Lang 2018; Fisher et al. 2018). However, this approach is prone to miss detection of beneficial mutations due to weak effects or genetic interactions (Buskirk, Peace, and Lang 2017). Unambiguously distinguishing beneficial mutations from hitchhikers requires measuring the fitness effect of all mutations within an individual evolved clone. Previously we systematically quantified the fitness effects of mutations in haploids using a bulk-segregant approach to generate large pools of segregants that contain random combinations of evolved mutations (Buskirk, Peace, and Lang 2017). Here we extend this approach to diploid populations.

Diploids adaptation differs from haploids in several important ways. First, diploids adapt more slowly compared to haploids (Marad, Buskirk, and Lang 2018; McDonald, Rice, and Desai 2016; Gerstein and Otto 2011; Orr and Otto 1994). This is because, in diploids, heterozygous recessive mutations are not acted on by selection (Orr and Otto 1994; Diamantis Sellis et al. 2016; Haldane 1924), and dominant beneficial mutations are comparatively rare (Deutschbauer et al. 2005; Zörgö et al. 2012). Evolved diploid genomes are, therefore, expected to contain fewer and/or weaker beneficial mutations. In addition, diploid genomes are expected to contain beneficial mutations and hitchhikers in both the heterozygous and homozygous states. Third, diploid populations are more likely than haploids to acquire aneuploidies and recessive deleterious mutations, both of which will interfere with the ability to genetically separate co-evolved mutations in evolved genomes. Distinguishing beneficial mutations from hitchhikers is, therefore, more challenging in diploid genomes.

Previously we sequenced 24 evolved diploid clones and identified adaptive mutations based on recurrence (Marad, Buskirk, and Lang 2018). To quantify the fitness effects of all mutations in a subset of these populations, we combined our previous bulk segregant approach with DNA barcoding (Buskirk, Peace, and Lang 2017; Ba et al. 2019). We find that while diploid clones carry ∼20-25 mutations each, only ∼6% of mutations are non-neutral with effects ranging between ∼1% and 5.4%. Beneficial mutations in haploids have larger effect sizes (1-10%) but are accompanied by half-as-many hitchhiking mutations (Buskirk, Peace, and Lang 2017). Allele reconstructions show that heterozygous adaptive mutations are frequently overdominant and homozygous adaptive mutations are either partially or fully dominant, consistent with Haldane’s sieve. These data argue that both smaller fitness effects and the reduced availability of beneficial mutations contribute to the slower adaptation in diploids. Finally, we show how changes to the environment impact the selective pressures in our experiment.

## RESULTS

### Diploid populations harbor recessive deleterious and recessive lethal mutations

According to Haldane’s sieve, recessive beneficial mutations fail to fix in asexual diploid populations because selection cannot act on the heterozygote (Connallon and Hall 2018; Haldane 1924). By similar logic, selection should be unable to prevent the accumulation of recessive deleterious mutations in asexual diploid populations. To test whether diploid populations contain recessive deleterious mutations we sporulated 17 diploid populations from Generation 4,000 and 9 clones from diploid populations from Generation 2,000 of a previously performed experimental evolution (Marad, Buskirk, and Lang 2018). All populations and clones produced tetrads; therefore, sporulation ability was maintained. Though sporulation efficiency was uniformly high, spore viability was not. At one extreme, Population C04 failed to produce a single viable spore across ten tetrads, indicating a severe meiotic defect, while five of the seventeen populations and two of the nine clones had spore viability at least as high as the ancestor (Table S1). However, even for populations and clones with high spore viability, the patterns of colony size segregation show that each of these contain at least one recessive deleterious mutation, and over half carry at least one recessive lethal mutation.

### Diploid evolved clones carry homozygous and heterozygous adaptive mutations

To identify beneficial mutations in diploid populations, we performed bulk-sporulation of evolved clones followed by bulk-mating to their ancestor to create large barcoded pools of diploid segregants that contain random combinations of heterozygous evolved mutations (Figure 1). To assess the ability of our methodology to recover maladaptive mutations, we pilot it on strains with recessive lethal mutations. Prior studies have shown that lethal haplotypes can be recovered as long as mating with a complementing strain is introduced soon after meiosis (Haarer et al. 2011). To show that we can recover recessive deleterious and recessive lethal mutations in our pools, we first performed a pilot experiment using six strains, five of which are hemizygous for an essential gene covering a diversity of biological processes: protein folding (*CNS1*), actin cytoskeleton (*ACT1*), cytokinesis (*IQG1*), karyogamy and spindle pole body formation (*KAR1*), and secretion (*SEC27*), and one which is hemizygous for a non-essential gene involved in N-glycosylation (*YUR1*) as a control (Figure S2). In all cases, we recovered the recessive-lethal mutation, though below the expected 1/2 rate (recovery ranged from 1/1000 for *sec27*Δ to 1/8 for *kar1*Δ, Figure S2).

**Figure 1.**
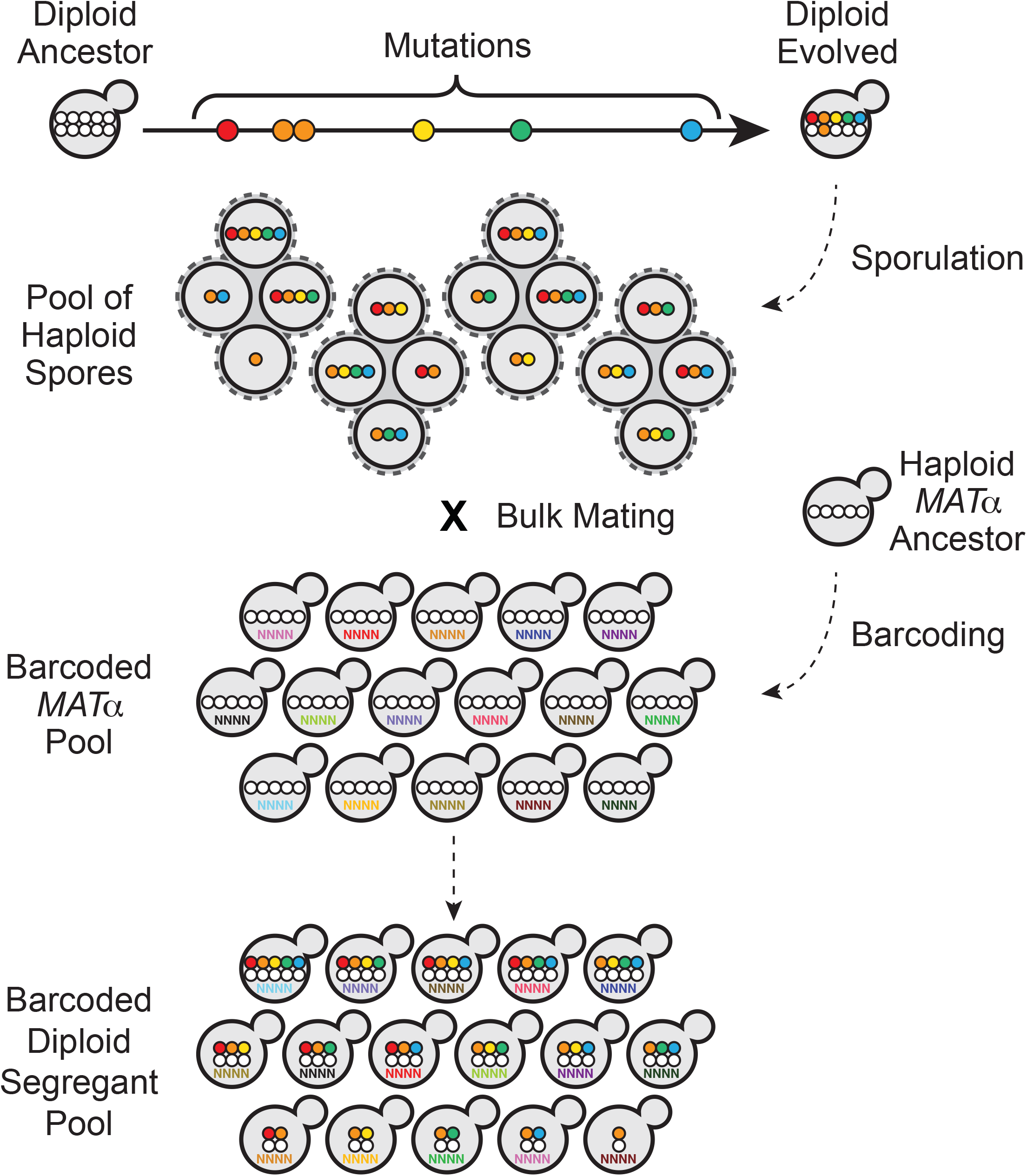
Cartoon of the strategy used to generate diploid segregant pools. Diploid evolved clones isolated from generation 2000 were sporulated and mated *en masse* to barcoded *MAT*α version of the ancestor. Each unique barcode represents a single mating event. The resulting diploids have genotypes intermediate to their ancestor and evolved parents and carry a barcode. Mutations homozygous in the evolved parent (in orange) end up in heterozygosity in all segregants. Figure S1 shows the barcoded locus.

To identify beneficial mutations in diploid evolved populations, we performed bulk-sporulations of six clones from Generation 2,000 followed by bulk-matings to create large barcoded pools of diploid segregants that contain random combinations of heterozygous mutations, each from an evolved diploid clone (Figure 1). Populations (A05, A07, F04, H06) were chosen on the basis of high fitness, high spore viability, and absence of aneuploidies in clones from the same generation (Marad, Buskirk, and Lang 2018). Because our strain background harbors the killer virus (a cytoplasmic dsRNA virus that can impact fitness (Buskirk, Rokes, and Lang 2020)), we assayed killing ability and toxin sensitivity for each clone and monitored killing ability throughout pool construction (data for parental clones shown in Table 1).

**Table 1.** Population and clone phenotypes.

| **Population** | **Population fitness\*** | **Pop. Killer activity\*\*** | **Clone** | **Spore Viability** | **Clone killing activity \*\*** |
| --- | --- | --- | --- | --- | --- |
| A05 | 5.47% | 0.67 | A05-C | 49.5% | 0.67 |
|  | - | - | A05-D | 50.0% | 0.33 |
| A07 | 5.60% | 0.67 | A07-C | 17.5% | 0.67 |
| H06 | 6.14% | 1 | H06-C | 51.0% | 1 |
| F04 | 6.41% | 1 | F04-C | 45.0% | 0.67 |
|  | - | - | F04-D | 92.5% | 1 |

\* Fitness values are from Marad et al. 2018
\*\* Score: 0 (no killing) to 1 (normal killing). No clones are sensitive to Killer toxin.

For each evolved clone, we generated two segregant pools: a “small pool” of ∼200 individuals and a “large pool” of >60,000 individuals. We used the small pools to quantify the segregant fitness via pooled competition assays and barcode sequencing (Venkataram et al. 2016) and the large pool to perform whole-genome whole-population sequencing. Barcoded derivatives of the ancestor and evolved clones were spiked into the segregant pools, which were then propagated under conditions identical to the evolution experiment for 110 generations in two replicates. In addition, we measured the fitness of the ancestor and evolved barcoded derivatives using a fluorescence-based fitness assay (Figure S3A). Fitness values from these independent assays are highly correlated (Figure S3B, R=0.92).

Homozygous and heterozygous beneficial mutations affect the segregant fitness distribution in different ways. Homozygous mutations in the evolved parent end up as heterozygous mutations in all segregants (Figure 1). Because homozygous beneficial mutations must be at least partially dominant to escape Haldane’s sieve, the fitness of segregants derived from evolved clones carrying such mutations is between those of the ancestor and the evolved parent. This pattern is observed in the segregants from clones A05-C, A05-D and A07-C (Figure 2, Table S2, and Extended Data Tables E1-E6). The fact that the distributions of fitness effects of the segregants from these clones is unimodal suggests that there are no heterozygous adaptive mutations in these clones.

**Figure 2.**
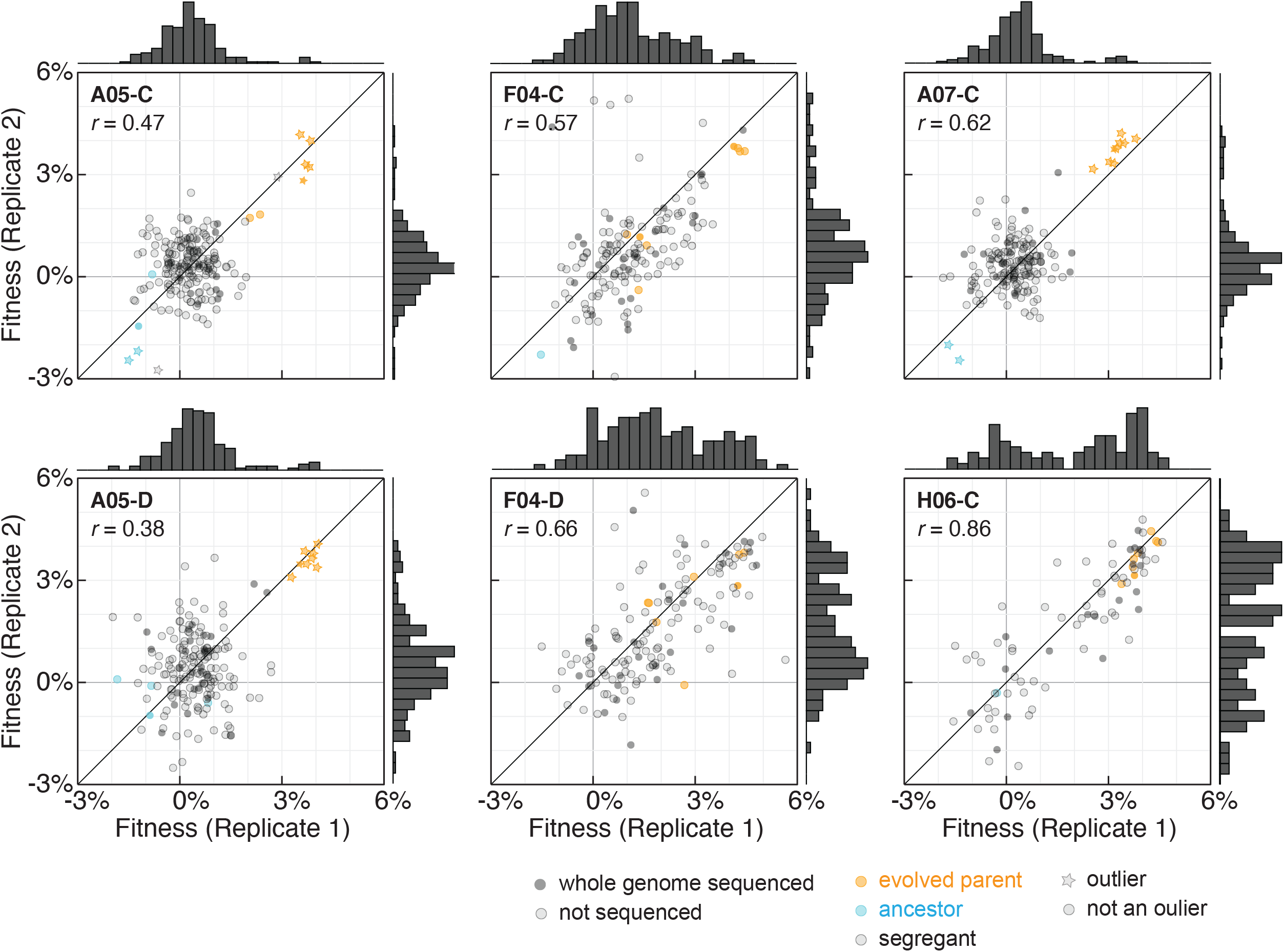
Fitness distributions of diploids derived from mating between a haploid version of the ancestor and the meiotic progeny of evolved diploid clones. The assays were barcode-based and the fitness was estimated with the algorithm published in (Venkataram et al. 2016). Fitness correlations are shown for 2 replicates per clone. Ancestor and evolved parents are annotated in cyan and orange, respectively. Derived diploids are annotated in grey. Clones for which there are whole genome sequencing data available are annotated with a closed circle. Outliers were identified via boxplot and Rosner tests performed in R (Tables S3 and Extended Data Tables E1-E6). Pearson correlations for each replicate pair is shown at the top left of each panel.

By contrast, the fitness distributions of the pools of segregants from the F04-C, F04-D, and H06-C evolved clones suggest fitness gains in these clones due to heterozygous adaptive mutations (Figure 2). The evolved and ancestral parents are not detected as outliers in these populations, but as expected for a non-transgressive segregation, they are at the extremes of the segregant distributions [Wilcoxon *p*-values: ancestral – segregants comparisons < 0.1 for individual replicates (ancestral fitness values were not resulted in both replicates), and segregants – evolved comparisons < 0.04 for averaged replicates].

To identify the homozygous beneficial mutations in A05-C, A05-D, and A07-C and the heterozygous beneficial mutations in F04-C, F04-D and H06-C, we sequenced the ancestral and evolved clones as well as 160 segregants across the six small pools. We genotyped each strain and identified the corresponding barcode sequence. The evolved clones carry on average 1 homozygous and 20 heterozygous mutations (all mutations shown in Table S3). Sequencing shows that both clones from population A05 have a single homozygous mutation in the common target of selection *ACE2*. Clone A07-C has three homozygous mutations, including in *WHI3* and *CTS1. CTS1* is the most common target across diploid populations and *CTS1* mutations are always observed in homozygosity due to their proximity to the highly recombinogenic rDNA locus (Marad, Buskirk, and Lang 2018). *WHI3* is not a known target of selection. The third homozygous mutation in A07-C is at an intergenic position on Chromosome II. Population H06 carries heterozygous mutations in common targets of selection *PDR5* and *PTR2*. Population F04 carries heterozygous nonsynonymous mutations in the common targets of selection *ACE2*, and *KRE6* (Marad, Buskirk, and Lang 2018). Interestingly, the two clones of F04 show subtly different behavior in all assays. Segregants from population F04-C show more of a continuum of fitness values between ancestral and evolved clones (Figure 2), with the evolved clone appearing as an outlier in one replicate. Furthermore, the genetic data suggest that F04-C, but not F04-D carries at least one recessive deleterious mutation (Table S1).

### Most genome evolution in laboratory-evolved diploids is non-adaptive

In addition to identifying heterozygous and homozygous beneficial mutations, we measured the fitness effects of all heterozygous mutations in each evolved clone using two methods. First, we combined the genotype information of a subset of segregants from whole genome sequencing and the fitness from barcode sequencing. Fitness effect of each variant was estimated as the difference in mean fitness of segregants with and without the variant (Figure 3, Table S4). To assess significance of the effect we used Bonferroni-corrected t-tests and Wilcoxon rank-sum tests (Extended Data Tables E7-E12). Few heterozygous mutations appeared significant in the evolved clones that harbored beneficial homozygous mutations, and none were significant after the Bonferroni correction (Extended Data Tables E7-E9). In populations with heterozygous beneficial mutations, we initially identified up to four putatively significant mutations (Figure 3). However, after correcting for co-segregation of mutations in our sequenced segregants (ANOVA, Extended Data Tables E10-E12), we show that there are two adaptive mutations in each clone (Table 2). Clones from population F04 carry heterozygous beneficial mutations in *ACE2* and *KRE6* with average fitness effects 1.33 % and 1.89%, respectively, for clone F04-C and 1.76% and 2.42%, respectively, for clone F04-D. Clone H06-C carries adaptive mutations in *PDR5* and *PTR2* with average fitness effects 3.57% and 2.03%, respectively.

**Table 2.** Adaptive mutations in diploids.

| Mutation | Clone | Evolved state | Segregant Fitness |  | Reconstruction Fitness |  |  |  |
| --- | --- | --- | --- | --- | --- | --- | --- | --- |
|  |  |  | Heterozygous | Homozygous | Heterozygous | Background | Homozygous | Background |
| *ace2*-R617G | A05-C | homozygous | 1.65% | 4.48% | ND | n/a | ND | n/a |
| - | A05-D | homozygous | 0.97% | 4.22% | - | - | - | - |
| *cts1*-G109D | A07-C | homozygous | 2.15% | 5.36% | ND | n/a | ND | n/a |
| *ace2*-R669* | F04-C | heterozygous | 1.33% | ND | 1.84% | evolved | 4.25% | evolved |
| - | F04-D | heterozygous | 1.76% | ND | - | - | - | - |
| *kre6*-S453L | F04-C | heterozygous | 1.89% | ND | 7.25% | ancestral | -11.28% | ancestral |
| - | - | - | - | - | 5.06% | evolved | 4.30% | evolved |
| - | F04-D | heterozygous | 2.42% | ND | - | - | - | - |
| *pdr5*-F562I | H06-C | heterozygous | 3.57% | ND | ND | n/a | ND | n/a |
| *ptr2*-V243fs | H06-C | heterozygous | 2.03% | ND | ND | n/a | ND | n/a |
| *pdr3*-Y276S | - | heterozygous | ND | ND | 4.81% | ancestral | -9.36% | ancestral |
| *pse1*-Q651* | - | heterozygous | ND | ND | 2.27% | evolved | -6.10% | evolved |
| *kre6*-A521T | - | heterozygous | ND | ND | 4.45% | evolved | ND | n/a |

**Figure 3.**
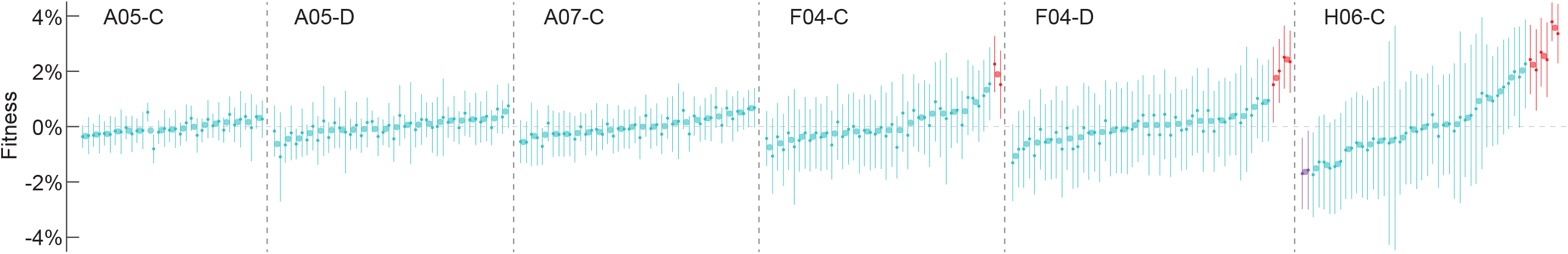
Quantification of heterozygous variant fitness effects using timecourse barcode sequencing and segregant WGS. Annotations reflect population and clone. Only mutations with data in both replicates are included. Mutations are ranked by mean average fitness for both replicates, where mean fitness was calculated as the difference evolved-ancestral allele mean fitness. Error bars represent 95% confidence intervals for replicate 1 on the left and replicate 2 on the right, of the t-test comparing evolved and ancestral alleles mean fitness. Mutations with neutral effect are considered those whose t-test confidence interval encompasses zero of at least one replicate and are annotated in light blue. Of the rest, mutations with positive or negative effects are annotated in red and purple, respectively. Data reported in Table S5 and more in-depth analysis in Tables Extended Data Tables E7-E12.

To corroborate these results, we also quantified the fitness effects of heterozygous mutations by propagating the big segregant pools (∼60,000 segregants each for clones A05-C, A05-D, A07-C, F04-C and F04-D) for ∼110 generations. As previously we estimated the fitness effect by tracking the frequency of each mutation by whole-genome sequencing every 20 generations with a minimum of 60x coverage (Buskirk, Peace, and Lang 2017). This data was noisy and contained a high rate of both false-positives and false-negatives (Tables S5, S6). Nevertheless, the *KRE6* mutation was identified in both clones from population F04, but in one replicate each and significance after correction holds only for the one clone. *ACE2* appeared significant in a single replicate with both methodologies but was not significant after correction.

Overall, we find that the majority of heterozygous mutations (∼96%) in our evolved clones are neutral, with the remaining ∼4% having modest fitness benefits ranging from 1.3% to 3.6% (Tables 2, 3). Among homozygous mutations, which make up ∼10% of the total evolved, the proportion of beneficial mutations is much higher (38%) and their fitness effects are larger (4.2% to 5.4%). Considering both heterozygous and homozygous mutations, ∼6% are adaptive in our evolved diploid clones. For comparison, in haploids ∼20% of mutations are beneficial with fitness effects up to 10% (Buskirk, Peace, and Lang 2017). These results are consistent with a slow rate of adaptation in diploids relative to haploids (Marad, Buskirk, and Lang 2018).

**Table 3.** Adaptive genome evolution in diploid and haploid populations.

| **Background** | **Mutations\*** | **Beneficial**<br>**Mutations\*** | **Adaptive**<br>**Evolution** | **Non-adaptive**<br>**Evolution** |
| --- | --- | --- | --- | --- |
| *Diploid* |  |  |  |  |
| Heterozygous | 139 | 6 | 4% | 96% |
| Homozygous | 8 | 3 | 38% | 62% |
| Total | 147 | 9 | 6% | 94% |
| *Haploid* |  |  |  |  |
| Total | 116 | 24 | 21% | 79% |

\*We considered each clone as an independent genotype, therefore mutations that appeared in both clones from the same population are counted twice. Otherwise, the number of hitchhikers would be artificially inflated relative to beneficial mutations.

### Heterozygous beneficial mutations are frequently overdominant

Short-term evolution in diploids is predicted to favor selection for overdominant mutations (Diamantis Sellis et al. 2016; D. Sellis et al. 2011). We determined the dominance of beneficial mutations using three approaches: forcing loss-of-heterozygosity in the evolved strains, backcrossing the evolved strains, or reconstructing the evolved mutations in the ancestral background. To force loss-of-heterozygosity in evolved diploid clones, we selected specific putative beneficial mutations from our previous work (Marad, Buskirk, and Lang 2018) based on three criteria. First, we selected mutations in common targets of selection. Second, we considered mutations in euploid populations (Marad, Buskirk, and Lang 2018). Third we avoided populations with linked heterozygous mutations to avoid losing heterozygosity at multiple loci (Fisher et al. 2018).

We used the CRISPR/Cas9 system to force gene conversion of individual heterozygous evolved alleles within evolved strains towards homozygosity for either the ancestral or evolved allele. We performed competitive fitness assays on these strains and the evolved heterozygote to ascertain the fitness effects of the mutations in the context of the evolved background (Figure 4A, Table 2). Three mutations (*bck1-*S945S, *bst1-*S740R, and *ubp12*-V279L) are neutral with no impact on the fitness of the evolved background when heterozygosity is lost in either direction (t-test, p > 0.3). Three mutations (*pse1*-Q6562*, *kre6*-S453L and *kre6-*A521T) are overdominant, with highest fitness while in heterozygosity (t-test, p < 0.05 for the *pse1*-Q6562* and *kre6-*A521T alleles and p = 0.057 for the *kre6*-S453L allele). *PSE1* was previously identified as a target in the autodiploid dataset, exclusively in heterozygosity (Fisher et al. 2018). Of those, we were unable to recover the homozygous mutant for *kre6-*A521T suggesting lethality. The last mutation (*ace2*-R669*) showed partial dominance. *ACE2* is a common target that is observed as both heterozygous and homozygous (Marad, Buskirk, and Lang 2018).

**Figure 4.**
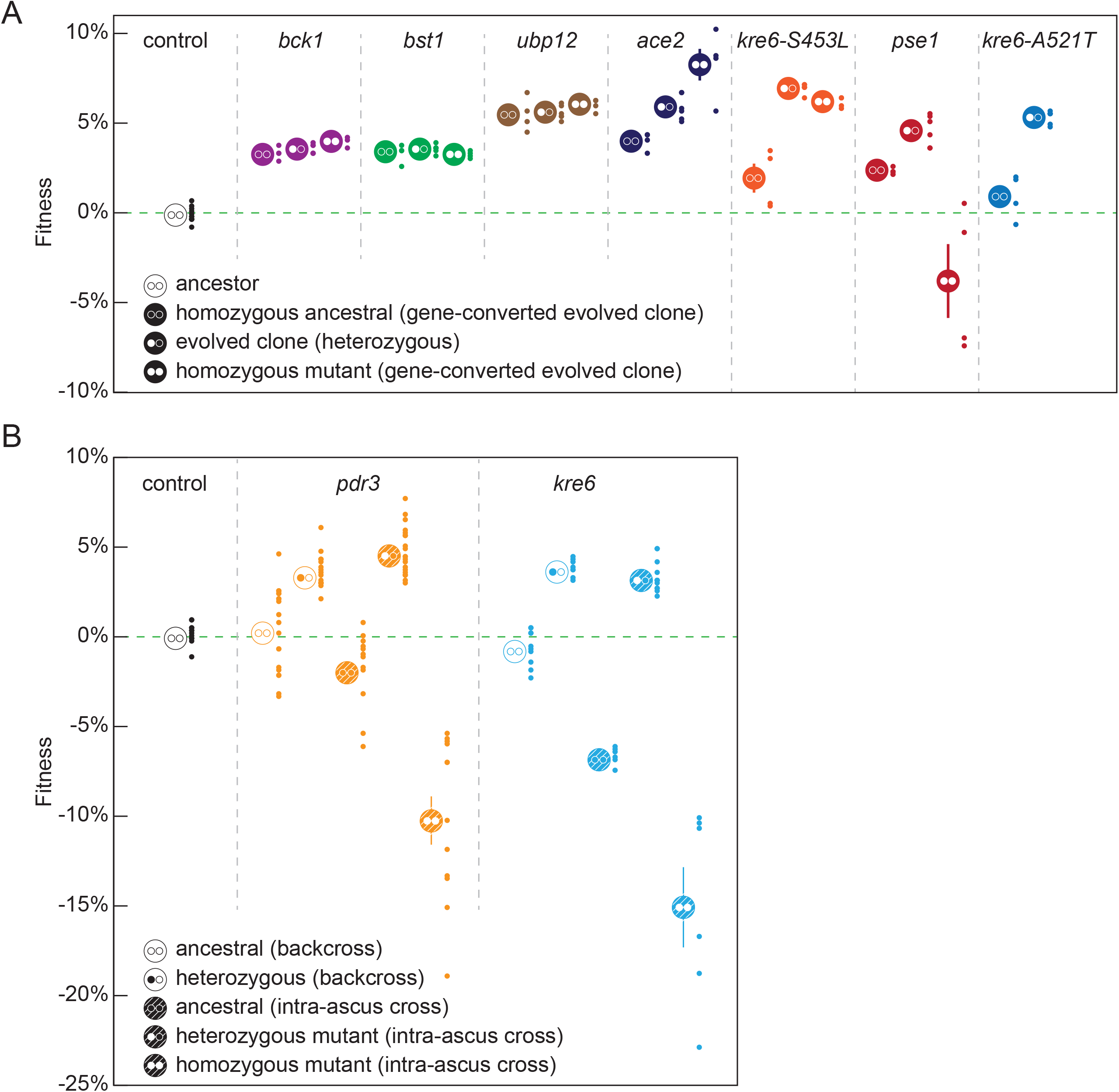
Overdominance of diploid-evolved heterozygous mutations. Average fitness of LOH of putative beneficial mutations was determined by driving LOH in the evolved background (**A**) or by introgressing the evolved mutations into the ancestral background (**B**). Average fitness is indicated by the large marker and small markers are the individual fitness values from independently constructed strains. The homozygous ancestral alleles are annotated by two open circles, heterozygous by one open and one white and homozygous evolved by two white circles. Open big circles in (B) annotate final segregant backcrossed to haploid version of the ancestor and filled circles annotate intra-ascus matings of the final cross.

Independently, we introgressed *pdr3-*Y276S and *kre6*-S453L in the ancestral background to generate heterozygotes and homozygotes. We chose *pdr3-*Y276S because this mutation co-segregated with a small colony phenotype (Table S1). For each mutation, we performed five rounds of backcrossing to replace most of the evolved background with the ancestral, while selecting for the allele of interest. Haploid segregants of the fifth round were used to construct diploids heterozygous and homozygous for the query allele, whose fitness was determined against a fluorescent version of the diploid ancestor (Figure 4B, Table 2). We find that both the *pdr3-*Y276S and the *kre6*-S453L mutants are beneficial (4.0% and 3.3%, respectively) if heterozygous, but strongly deleterious (−10.2% and −15.0%, respectively) if homozygous (Figure 4B). Though both the forced loss-of-heterozygosity experiment and the backcrossing experiment show that the *kre6*-S453L mutation is overdominant, the deleterious effect of the homozygous mutation is far less severe in the evolved background (compare Figure 4A to Figure 4B).

### Adaptive mutations are pleiotropic

Despite the differences between the haploids and diploids in the rate of adaptation and distribution of fitness and dominance effects of beneficial mutations, the genetic targets of selection (given an otherwise identical evolutionary environment) impinge on similar biological processes, such as cell wall metabolism, drug transport, as well as nutrient sensing and signaling. We sought to identify these common selective pressures across ploidy states. Earlier studies on adaptation in the same experimental system have hinted on spatiotemporal heterogeneity providing the selective forces driving adaptation (Lang, Botstein, and Desai 2011; Frenkel et al. 2015). This heterogeneity arises from the fact that there is no agitation during growth leading to nutrient and oxygen gradients. Additional support is provided by evolution experiments showing that cell wall mutations impact rates of settling (William C. Ratcliff et al. 2015; Koschwanez, Foster, and Murray 2013).

In order to study selection in our environment, we measured the fitness of evolved clones H06-C and F04-D, as well as a reconstructed heterozygous *kre6-*S453L strain, in modified environments by removing antibiotics (which were included to prevent bacterial contamination (Lang, Botstein, and Desai 2011)) and/or agitating the 96-well plates during growth (Figure 5A). Clone H06-C displayed a strong trade-off on the well-mixed environments, in which it had an almost 5% fitness deficit compared to its ancestor, while it was completely unfazed by the antibiotics in the growth medium. Clone F04-D is always more fit than its ancestor, but the fitness advantage was decreased without antibiotics and/or with agitation. Interestingly, the heterozygous *kre6-*S453L strain and the F04-D clone (which contains this mutation) had the lowest fitness in the well-mixed environment with antibiotics, suggesting that antibiotics were inhibiting the growth of these mutants in the well-mixed environment.

**Figure 5.**
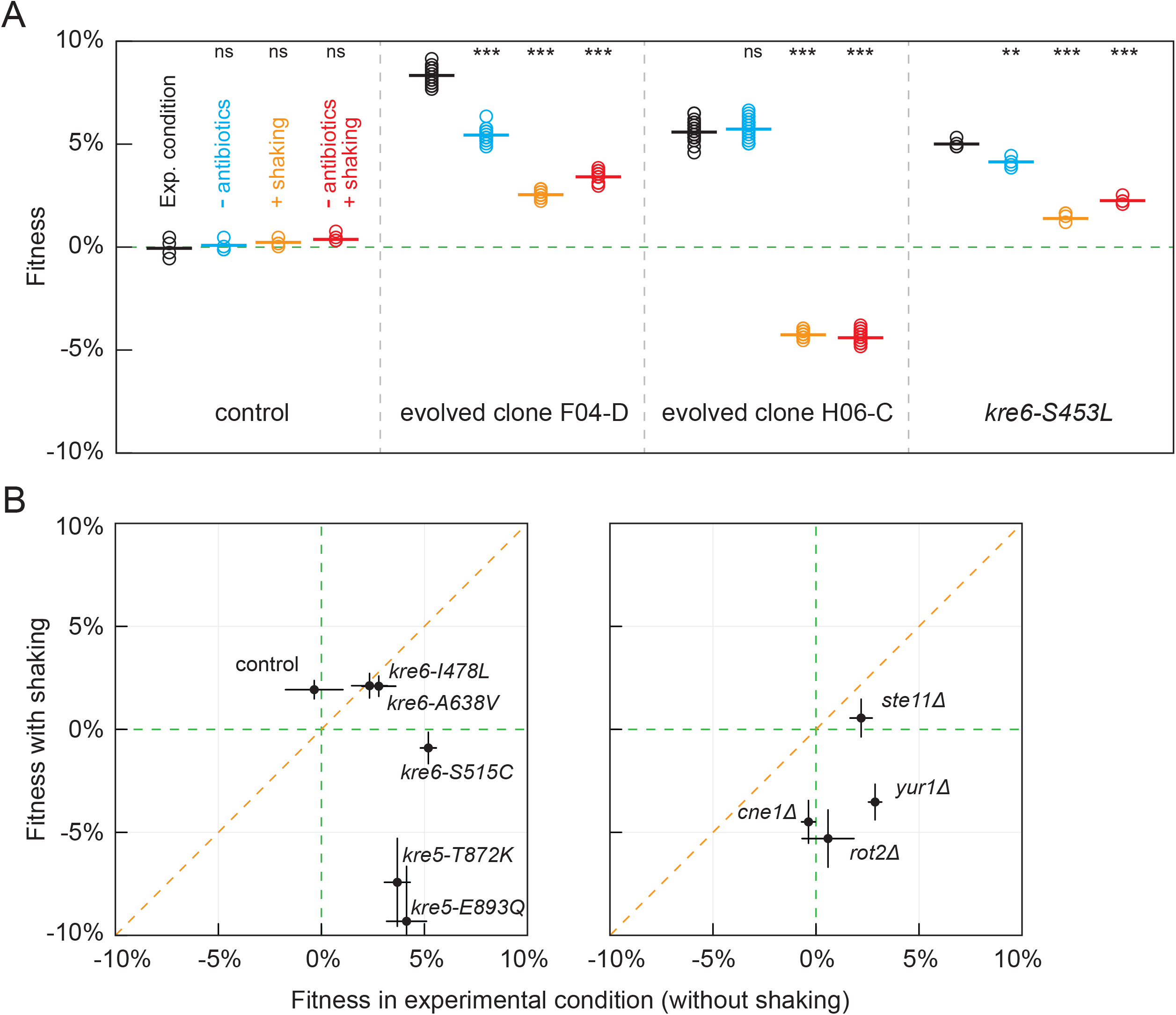
Fitness of evolved mutations changes upon environmental perturbation. Relative fitness was estimated against a fluorescent version of the respective ancestor. **A**. Evolved clones, H06-C and F04-D, fitness and the fitness of *KRE6* allele from population F04 engineered into the ancestral background were assayed in the different conditions as annotated on the 1^st^ panel. T-test significance is with respect to the experimental condition (ns: not significant, *: p<0.05, **: p<0.001, ***: p<0.0001). **B**. Mutations that emerged during haploid evolution (left panel) or loss-of-function mutations of genes mutated during haploid evolution (right panel) were engineered in the ancestral background and fitness was assayed in a static (evolutionary) and a well-mixed environment, shown on the x- and y-axis, respectively. Error bars represent standard error from at least 3 replicate assays.

To determine if this was a general effect of cell-wall mutations, we measured the fitness of a panel of cell-wall mutations from our haploids and specific gene deletions, in static and well-mixed environments with antibiotics (Figure 5B). *KRE6* and *KRE5* mutations showed a range of tradeoffs, with *KRE5* mutants being the most compromising in the well-mixed environment (Figure 5B). We also tested the fitness effects of loss-of-function mutations for *STE11, YUR1, CNE1*, and *ROT2. CNE1* and *ROT2* deletions did not phenocopy the evolved alleles, as they displayed no fitness in the evolutionary condition. The *STE11* and *YUR1* deletions displayed a fitness advantage in the static environment, but only *YUR1* manifested a strong trade-off in the well-mixed environment. Collectively, these data suggest that adaptation in our environment that relies on modifications of the cell wall is typically lost under well-mixed environments.

To assess the impact of environment on all mutations, we used fitness estimates of the genotyped barcoded segregants in the different environments (Figure S4, Tables S7-S10, Extended Tables E11 and E12). We found that removing antibiotics causes a wider spread in the fitness distribution (Figure S4). The evolved parent H06-C displayed strong trade-offs in the well-mixed environments, but only a few segregants appeared to approach the fitness deficits of the evolved parent, suggesting a multi-allelic effect. Additionally, *PTR2* is adaptive in all environments, but *PDR5* is adaptive only in static environments. In addition, a mutation in *FLO9* appears to have a small fitness deficit in the well-mixed environments (−0.7% in 3 out of the 4 assays) and a variant in *ASK10* a small fitness advantage in the mixed environment without antibiotics. Assays in segregants from clone F04-D supported the earlier seen advantage of *KRE6* and *ACE2* mutations in only one replicate of the evolutionary condition, presumably because of the scarcity of information in this round. Unexpectedly, *ACE2* had a positive effect ∼1% in the well-mixed environment without antibiotics and in one replicate in the well-mixed environment with antibiotics. *KRE6* did not appear as consequential in any other environment. Three more mutations (a missense in *SNU114*, a missense in *VPS41* and an intergenic mutation) had a significant deficit in a single environment each (in well-mixed no antibiotic, in well-mixed with antibiotic and in the evolutionary condition, respectively) but only in one of the two replicates.

## DISCUSSION

We quantified the fitness effects and the degree of dominance of mutations that emerged during diploid evolution using a combination of bulk-segregant fitness assays, genetic reconstructions and backcrossing. Collectively four principles emerge from this study. First, diploid populations harbor fewer driver mutations and more hitchhiker mutations relative to haploid populations evolved in the same condition. Second, diploid populations accumulate many recessive deleterious or lethal mutations that reduce spore viability. Third, all beneficial mutations in diploids display some degree of dominance on their fitness effect. Fourth, most heterozygous beneficial mutations are overdominant.

We find that short-term clonal evolution in diploids is largely non-adaptive, as only 6% of the identified mutations contribute to fitness gains. We identify both heterozygous and homozygous beneficial mutations. Consistent with our work in haploids we did not identify any beneficial synonymous or intergenic mutations. Diploid populations also carry a load of recessive-deleterious and recessive-lethal mutations. In contrast, 20% of mutations in haploid populations were shown to be beneficial and less than 1% deleterious (Buskirk, Peace, and Lang 2017). This difference is also reflected by the fitness increase of the two ploidies over time: diploid fitness increased by an average of 5.8% over 4000 generation, whereas haploid fitness increased by 8.5% over 1000 generations (Marad, Buskirk, and Lang 2018; Buskirk, Rokes, and Lang 2019).

Our results support theoretical predictions and prior experimental results suggesting that heterozygous beneficial mutations are likely to be overdominant (Haldane 1924; D. Sellis et al. 2011; Diamantis Sellis et al. 2016). Even those mutations that are not overdominant (like *ace2* in population F04) are at least partially dominant. We previously observed the inverse pattern for beneficial mutations that evolved in haploid populations: they are either recessive or underdominant (Buskirk, Peace, and Lang 2017; Marad, Buskirk, and Lang 2018; Fisher et al. 2018). Using data from our diploid, autodiploid and haploid datasets (Lang et al. 2013; Buskirk, Peace, and Lang 2017; Marad, Buskirk, and Lang 2018; Fisher et al. 2018), we summarize the patterns of dominance for haploid and diploid laboratory evolution experiments (Figure 6). Beneficial mutations that arose on the haploid and diploid backgrounds occupy separate but overlapping regions of this space. Note that the same genes often underlie adaptation in both haploids and diploids, but their dominance effects are different depending on the background in which they arose.

**Figure 6.**
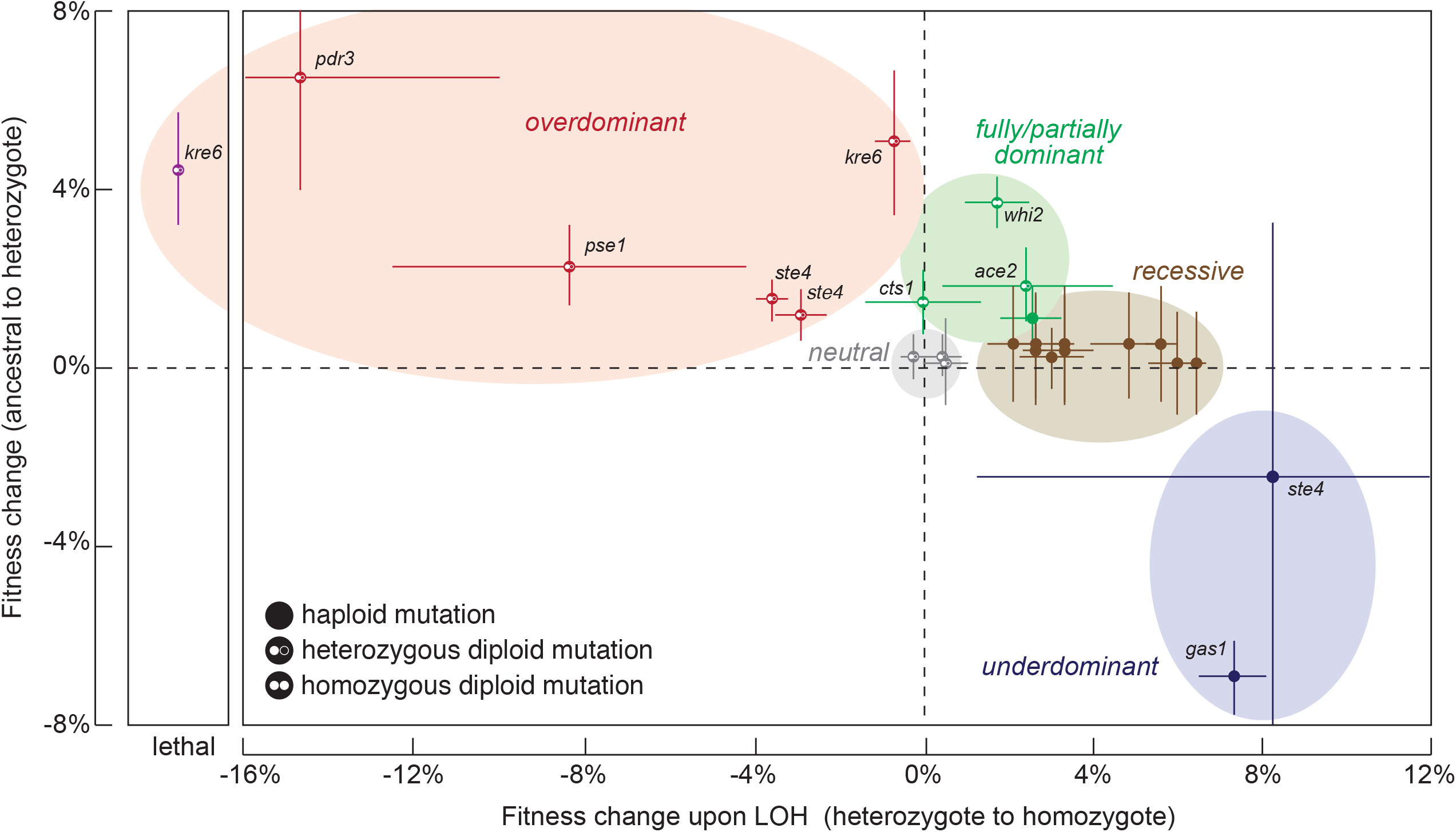
The degree of dominance of beneficial mutations is dependent on the ploidy of the background on which they arose. The fitness effect of beneficial mutations from diploids, autodiploid, and haploid datasets (Fisher 2018, Marad 2018, Buskirk 2017, Lang 2013) are shown on a single plot. The background on which each mutation arose is indicated by the symbol, and select genes are identified by name. Overdominant mutations are in the orange-shaded region and arise exclusively as heterozygous mutations in diploids. Only diploids have access to these mutations. Partially dominant mutations are in the green-shaded region. Both haploids and diploids have access to these mutations. Recessive mutations and underdominant mutations are in the brown-shaded and purple-shaded regions, respectively, and arise exclusively in haploid backgrounds.

We show that a bulk-segregant approach can be used to identify and quantify the fitness effects of all heterozygous mutations in evolved diploid populations. By mating immediately after breaking the tetrad asci, we show that our method can partially recover recessive lethal mutations in the final pools. In addition, by including the ancestor and evolved diploid parents into the segregant pools, we are able to identify populations with homozygous beneficial mutation and quantify their combined fitness effects. This is a useful feature of our assay because, although homozygous mutations contributed only ∼5% of the total genetic variation, they contributed ∼33% of the adaptive variation (Table 3).

By altering specific environmental parameters, we uncovered one condition that appears to exert a dominant selective pressure, driving adaptation in our system. Populations growing statically in liquid media, as is the case in our environment, are subjected to nutrient and oxygen gradients, as well as the spatial structure imposed by the shape of the well, which has been previously shown to drive evolutionary dynamics and adaptations in that same and similar systems (Frenkel et al. 2015; W. C. Ratcliff et al. 2012). When adapted clones were exposed to a well-mixed environment, they lost competitive advantage with respect to their ancestor. However, the specific patterns of the evolved parents and their segregants’ fitness effects across environments suggest complex origins of the trade-offs, due only in part to the adaptive mutations. Trade-offs have been observed before upon reversion of the selective pressure that drives adaptation in a system (Hietpas et al. 2013). Adaptive responses manifesting trade-offs have been observed in the loss of signaling/regulatory pathways, which is adaptive in constant environments (chemostat) but becomes maladaptive in oscillating environments (batch culture) (Kvitek and Sherlock 2013), and in adaptations driven by carbon limitation resulting in fitness costs when carbon is in excess (Wenger et al. 2011).

We quantified the fitness effects of all mutations in six diploid clones from four 2,000-generation evolved populations. We show that ploidy has a large impact on genome-sequence evolution, selecting for partially dominant and overdominant beneficial mutations, but allowing for the fixation of a higher proportion of neutral and recessive-deleterious genetic hitchhikers. Over the short-term, overdominance will lead to the maintenance of heterozygosity of linked loci (Fisher, Vignogna, and Lang 2021). Over long timescales, however, it is expected that modifiers arise to alleviate the negative effects of these overdominant mutations (Diamantis Sellis et al. 2016; D. Sellis et al. 2011).

## MATERIALS AND METHODS

### Yeast strains and strain construction

The strains used in this experiment are derived from diploid yGIL672 (W303 background), with genotype *MAT***a**/α, *ade2-1, CAN1, his3-11,15, leu2-3,112, trp1-1, bar1*Δ::*ADE2, hml*αΔ::*LEU2, GPA1*::*NatMX, ura3*Δ::p*FUS1*-yEVenus.

Evolved clones of yGIL672 were derived from generation 2,000 of a 5,000-generation experiment described previously (Marad, Buskirk, and Lang 2018). Killing ability and sensitivity to killer toxin were assayed as described previously (Buskirk, Rokes, and Lang 2020) using a modified version of the standard halo assay. Killing ability was assayed against a hypersensitive tester strain (yGIL1097) and killer toxin sensitivity was assayed against yGIL432.

Gene deletions were introduced by amplifying the KanMX cassette from the hemizygous or *MAT***a** deletion collections (Euroscarf) and transforming using the standard lithium acetate protocol (Gietz and Schiestl 2007). Lethality of the essential gene deletions was verified via tetrad dissection. Evolved mutations were introduced into the ancestral background (yGIL432, yGIL646, and yGIL672; *MAT***a**, *MAT*α, and *MAT***a**/α respectively) using CRISPR/Cas9 allele swaps as described previously (Fisher, Kryazhimskiy, and Lang 2019). Briefly, oligonucleotides specifying the gRNA were hybridized and introduced into the *SWA1* and *BCL1* restriction sites of pML104 (Addgene #67638). Repair templates containing evolved mutations were generated by amplifying ∼500 bp fragments centered around the mutation of interest or by gBlock synthesis (IDT) containing the mutation or interest along with synonymous PAM site changes. For driving loss-of-heterozygosity (LOH) in diploids, no repair template was provided. All plasmids and strain constructions were validated by Sanger Sequencing (Genewiz or Psomagen).

### Barcode library transformation

The low complexity barcode was amplified out of a yeast library [strain XLY092 (Liu et al. 2019)] in 3 overlapping amplicons 1-2 kb long and the amplicons were introduced into yGIL432 via a lithium acetate transformation (Gietz and Schiestl 2007). The amplicons were targeted upstream of the *GPA1* locus (primers in Table S11), replacing the NatMX cassette that marks our ancestors and all derivatives. The choice of *GPA1* locus has the advantage of being located close to chromosome VIII centromere, thus reducing the chance of gene conversion, and LOH. Successful transformants were selected on YPD supplemented with kanamycin. Integration at the correct locus was screened via lack of growth on media supplemented with CloNAT and amplification of the expected junctions and/or receptiveness to the high complexity barcode [pBAR7-L1 plasmid library (Liu et al. 2019)]. The high complexity barcode library was introduced to low complexity individual transformants via a following transformation. Following heat shock Cre recombinase was induced overnight in liquid YP + 2% galactose. Transformants were selected on synthetic complete media lacking uracil and harvested via pooling after 2 days.

### Construction of barcoded bulk segregant pools

Sporulating cultures of evolved diploid clones were mated each with a different pool of the barcoded strain with an estimated diversity of 400,000 barcodes, as follows. 4 mL sporulating cultures (∼2×10^7^ tetrads/mL and ∼30% sporulation efficiency) were resuspended in 60 uL water and 2000 U zymolase (USBiological) and the samples were incubated at 30°C for 1 hr. Then 20 uL glass beads and 100 uL Triton X-100 were added, the samples were vortexed for 2 minutes, incubated at 30°C for 40 minutes and vortexed for an additional 2 minutes. 1.8 mL water was added, and the samples were sonicated at power 4 for 4 seconds. The solution was mixed with the haploid mating partner on nylon membranes (GVS, Sanford NE) at an excess of 1:100 (100 barcoded cells for each *MATα* spore) using a vacuum manifold. The membranes were incubated on a YPD plate at room temperature overnight and subsequently the cells were harvested from the membranes with PBS and were plated on synthetic complete medium lacking uracil and supplemented with CloNAT (selection for mated diploids) at different densities. Small pools were generated by picking 192 individual colonies off of each low-density CSM-ura+ClonNAT plate, propagating them in 96-well plates and mixing them at equal volumes. Large pools were generated by scraping ∼60,000 colonies off of high-density CSM-ura+ClonNAT plates. Control barcoded parental strains were derived from yGIL672. At each step the individual clones were isolated and assayed for fitness, killing ability and sensitivity to killer toxin. With the exception of the clones from population F04, phenotypes of the parental clones remained consistent throughout the barcoding process.

### Quantifying the recovery of recessive lethal mutations

To estimate mating efficiency of query haplotypes, sporulating cultures of diploid lab W303 derivatives, engineered with the respective gene deletions, were digested with zymolase (USBiological), the asci were broken, and the resulting spores were mass mated to a haploid mating partner, as follows: 3 mL sporulating cultures were resuspended in 60 uL water and 2000 U zymolase and the samples were incubated at 30°C for 1 hr. Then 20 uL glass beads and 20 uL Triton X-100 were added, the samples were vortexed for 2 minutes, incubated at 30°C for 40 minutes and vortexed for an additional 2 minutes. 1.2 mL water was added, and the samples were sonicated at power 4 for 4 seconds. The broken asci mix was mixed with the haploid mating partner either in patches on YPD plates or in membranes using a vacuum manifold. The resulting query diploids were quantified by plating to YPD supplemented with G418 and cloNAT. Percent recovery was estimated as the fraction of colony-forming units (cfu) on YPD media supplemented with G418 and cloNAT over the cfu on YPD supplemented with G418 (marker of the limiting mating partner) and expressed as a percentage of the non-essential control (*yur1Δ*) mating efficiency. For the mating efficiency calculation, it was assumed that sporulation efficiency was the same for all genotypes. Euploidy of the resulting query diploids was verified via tetrad dissection.

### Backcrossing of evolved clones

A backcrossing approach was used to introgress putative recessive lethal mutations into the ancestral background. We transformed evolved clones with plasmids carrying ancestral versions of genes harboring putative recessive lethal mutations (MoBY collection(Ho et al. 2009)). We sporulated each evolved clone and screened a single four-spore tetrad for the allele of interest. An appropriate spore was selected and backcrossed to either yGIL432 (*MAT***a**) or yGIL646 (*MAT*α). For each clone, we performed at least five backcrosses in order to remove other evolved mutations from these strains. Haploid progeny of the final tetrad dissection and haploid ancestral strains were used to generate a heterozygous backcrossed evolved mutation (cross of a segregant with evolved allele and the ancestor), a heterozygous intra-tetrad mating (cross of segregants with and without the evolved allele), a homozygous ancestral backcross (cross of a segregant without the evolved allele and the ancestor), a homozygous ancestral intra-tetrad mating (cross between two segregants with ancestral allele), and a homozygous mutant intra-tetrad mating (cross between two segregants with evolved allele). Each of these five types of crosses were performed in at least triplicate and the final strains were plated on 5-FOA to select for loss of the MoBY plasmid.

### Fitness Assays

Unless otherwise stated, all fitness assays were performed under the same conditions as the evolution experiment. The populations were diluted daily at 1:2^10^ using a BiomekFX liquid handler into 128 mL of YPD plus 100 mg/mL ampicillin and 25 mg/mL tetracycline to prevent bacterial contamination. The cultures were incubated at 30°C in an unshaken 96-well plate.

The fluorescence-based fitness assays were previously described (Buskirk, Peace, and Lang 2017). Briefly, the query strain was mixed isostoichiometrically with a fluorescent version of the ancestor (ymCitrine-labeled). The assays were performed for 30-50 generations and sampled every 10 generations (4-6 timepoints total). Following each transfer, 4 μL of the saturated culture were transferred in 60 uL PBS and the samples were stored at 4 °C for 1-2 days before being assayed by flow cytometry (BD FACSCanto II). Data were analyzed in FlowJo. Fitness was calculated as the linear regression of the log ratio of experimental-to-reference frequencies over time in generations.

For the barcode-based fitness assays, the small pools of 192 barcoded diploid segregants were mixed with four independently barcoded ancestral derivatives and between seven and eleven independently barcoded evolved derivatives (with volumes adjustments to achieve equal strain representation). Each pool was used to seed 2 columns of a 96-well plate (16 wells) and were propagated for 110 generations in the same way as the original evolution experiment, with sampling every 10 generations. For each timepoint, each column (8 wells) was pooled, the cells were spun down and stored at −20 °C for genomic DNA preparations. To monitor changes in allele frequencies, fitness assays were performed in the same way, using the large pools of ∼60,000 segregants.

### Library preparations and sequencing

#### Barcode determination of individual segregants

To identify the barcodes of the isolated segregants in 96-well plates, we employed a 3-(column, row, plate) dimensional pooling strategy, inspired by (Baym et al. 2016). Briefly, we inoculated 13 96-deep well plates with YPD from our frozen stocks (2 plates per mating and one control). The cultures grew for 2 days at 30°C and then the contents of wells were pooled by column, row and plate, resulting in 12, 8 and 13 pools, respectively. Genomic DNA was isolated and barcode libraries were prepared from the pools as described below.

#### gDNA preparation protocol

Cells from ∼1.5-2 mL saturated culture were resuspended in 100 µL lysis buffer (0.9 M Sorbitol, 50 mM Na Phosphate pH 7.5, 240 µg/mL zymolase, 14 mM ß-mercaptoethanol) and incubated at 37°C for 30 minutes. 10 µL 0.5M EDTA and 10 µL 10% SDS were added consecutively, with brief vortexing after each addition, and the samples were incubated at 65°C for 30 minutes and then on ice for 5’. 50 µL of 5 M potassium acetate were added, the samples were mixed, incubated on ice for 30 minutes and spun down at full speed in a microcentrifuge for 10 minutes. The supernatant was transferred to a new tube with 200 µL isopropanol and was incubated on ice for 5 minutes. The nucleic acid was spun down full speed in a microcentrifuge for 10 minutes and washed twice with 70% ethanol, was let to dry completely and then resuspended in 20 ul 10 mM Tris pH 7.5. Overnight incubation at room temperature or short incubation at 65°C sometimes was necessary for complete resuspension. RNA was digested with the addition of 0.5 µL 20 mg/ml RNase A (ThermoFisher Scientific, Waltham MA) and incubation at 37°C for 1 hour or at room temperature overnight.

#### Barcode sequencing libraries protocol

A two-step PCR protocol was used to amplify the barcoded locus (primers in Table S11). For the first amplification a maximum of 200 ng genomic DNA (corresponding to 7.5×10^6^ diploid *S. cerevisiae* genomes) was used as template in a 20 uL reaction with the following composition: 20 nM each forward and reverse primer (PU1 and PU2, Table S11), 10 ng/µL gDNA, 1 mM dNTPs, 0.2 uL Herculase II fusion DNA polymerase (Agilent, Santa Clara CA), 1X Herculase buffer, in the following conditions: hot start, initial denaturation at 98°C for 2’, 2 cycles of 98°C for 10”, 61°C for 20” and 72°C for 30”, and final extension at 72°C for 1’. Primers PU1 and PU2 introduce unique molecular identifiers and this first-step reaction is used as is as a template for the second step reaction, which introduces library-specific indexes for multiplex sequencing. To the first-step reaction, 30 uL with the following composition are added: 0.4 µL Herculase, 1xHerculase buffer, 1 mM dNTPs and 417 nM each of BC_i5 and BC_i7 (Table S11), and was amplified in the following conditions: hot start, initial denaturation at 98°C for 2’, 22 cycles of 98°C for 10”, 61°C for 20” and 72°C for 30”, and final extension at 72°C for 1’. DNA from all libraries was pooled isostoichiometrically, based on DNA concentrations estimated on Nanodrop. A 350 bp band was gel-purified of the final pool with the QIAGEN gel extraction kit (QIAGEN, Germantown MD).

#### Whole genome and whole genome whole population sequencing protocol

Genomic DNA was prepared for each of the segregants, after they were grown to saturation on YPD for whole genome sequencing. Whole-genome whole-population time-course sequencing of the fitness assays was performed to monitor changes in allele frequencies. Samples were thawed from - 20°C and sequencing libraries were prepared according to (Baym et al. 2015) with the modifications described in (Buskirk, Peace, and Lang 2017). Individual libraries were quantified by Nanodrop and pooled. Gel extraction included fragments in the 350-650 bp range.

#### Library QC and sequencing

Pooled samples were analyzed by BioAnalyzer on a High-Sensitivity DNA Chip (BioAnalyzer 2100, Agilent), before sequencing on an Illumina HiSeq 2500 sequencer with 250-bp single-end reads or on a NovaSeq with 2×150 bp paired-end reads at the Sequencing Core Facility within the Lewis-Sigler Institute for Integrative Genomics at Princeton University.

### Data analysis

Raw sequencing data were split by index using a dual-index barcode splitter (barcode_splitter.py) from L. Parsons (Princeton University).

#### Fitness estimation from barcode sequencing

Lineage fitness estimation from barcode sequencing data was performed as in (Venkataram et al. 2016). Raw barcode counts were prepared from barcode sequencing reads with use of existing software and a custom python script. Briefly, reads derived from paired-end sequencing were merged with pear (v0.9.11) (Zhang et al. 2014). Merged reads (or reads derived from single-end sequencing) were aligned against the expected barcoded locus sequence with bowtie2 (v2.3.4.1) (Langmead and Salzberg 2012). Barcodes and unique molecular identifiers (UMI) were extracted from the aligned reads and clustered using bartender (v1.1) (Zhao et al. 2018). Barcodes from reads were updated using the cluster centers derived from bartender and Levenshtein distance with threshold 2. The updated reads and the UMI were used to derive raw barcode counts, which were used as input to the lineage fitness algorithm.

#### Whole genome sequencing (WGS) analysis

Data from libraries from genomic DNA were subsequently trimmed from adaptor sequences using fastx_clipper from the FASTX Toolkit (http://hannonlab.cshl.edu/fastx_toolkit/download.html), version 0.0.14 if they originated from a single-end or trimmomatic, version 0.36 (Bolger, Lohse, and Usadel 2014) with option PE if they originated from a paired-end sequencing run. Each sample was aligned to the complete and annotated W303 genome using Burrows–Wheeler Aligner (BWA, v.0.7.15) (H. Li and Durbin 2009), option mem. Bam files were generated from sam files, sorted and indexed with samtools, version 1.4 (Heng Li et al. 2009). Bam files from libraries originating from the same sample were merged prior to sorting and indexing. Variants were called using FreeBayes, version 1.1.0 (https://github.com/ekg/freebayes), with option pooled-continuous for population data or option pooled-discrete with ploidy 2 for clonal data. VCF files were annotated using SnpEff, version 4.3 (Cingolani et al. 2012).

#### Variant discovery in evolved clones

We generated a consensus evolved variant list for each of the evolved parents, considering segregant, parental and population WGS data. First, we merged bam files derived from segregant libraries by common descend and from population libraries from the same initial pool. Merged segregant datasets consisted of 21 sequenced derivatives from clone A05-C, 28 from clone A05-D, 26 from clone A07-C, 24 from clone H06-C and 35 each from clones F04-C and F04-D. Merged population datasets consisted of 3 fitness assays each for initial evolved clones A05-C and F04-D and 2 assays each for initial evolved clones A05-D, A07-C, and F04-C. Each assay is made up of 6 timepoints. Downstream analysis was performed as described for WGS analysis. Variants were called with freebayes setting the ploidy option at 8 (considering that heterozygous mutations will appear in half the diploid derivatives in heterozygosities and trying to correct for small sample sizes. Mutations from the population merged datasets were called with parameters -F 0.01 -C 5 and --pooled-continuous. Parental variants parameters are as described for clonal data in WGS analysis. Each of these call sets were filtered as follows: calls that mapped on 2-micron plasmid or mitochondria, calls with low quality score (<19.99) and calls with more than one alternative allele were excluded. Additionally, total coverage of variant, fraction of alternative calls, as well as forward and reverse fraction of alternative calls were considered. In particular, we included calls with coverage z-score between −0.5 and 3 and forward to reverse alternative allele ratio between 0.4 and 2.5. The filtered variant list was then manually curated by visual inspection of the alignments on IGV (Robinson et al. 2011). We also computationally filtered the list using the following criteria for inclusion: The variant is called in at least 2 different datasets, one of which is the WGS of the evolved parent. The variant is called in a single population and not in the ancestor. The list that resulted after application of these criteria overlapped with the manually curated list that resulted after visual inspection of alignments. To specifically discover homozygous mutations, we applied the following 2 criteria for consideration in each of the 3 datasets: total coverage >29 and forward and reverse representation of the alternative allele. Subsequently we categorized mutations as heterozygous in the dataset (if alternative allele to total coverage was between 0.3 and 0.7) or homozygous if alternative allele to total coverage was greater than 0.7). Mutations that passed this filtering had to be called ‘homozygous’ in the parental dataset and ‘heterozygous’ in at least one of the merged segregants or merged populations datasets in order to be categorized as homozygous in the evolved parent.

#### *Assign fitness values to* mutations *from WGS and timecourse barcode sequencing of segregants*

Freebayes parameters for variant calling from clonal data were as described in WGS analysis. Evolved mutations in the consensus list were scored for presence/ absence in each segregant. Fitness values were attached to each genotype by using barcode and well coordinate information. For mutations represented by at least 3 fitness values in each of the presence, absence groups we estimated their fitness effect as the difference between the averages of the presence and absence groups. Significance was initially assessed by t-test and rank sum test and was Bonferroni-adjusted. Additionally, we performed ANOVA with input all mutations that appeared significant in at least one test before Bonferroni correction in at least one assay (including assays in the evolutionary condition and in conditions that deviate from the evolutionary when available).

#### *Assign fitness values to* mutations *from timecourse whole population WGS data*

Freebayes parameters for variant calling from individual timepoints data were -F 0.05 -C 3 and -- pooled-continuous. For each of the mutations in the consensus lists we calculated the natural logarithm of 2*evo/(anc-evo) per replicate assay and timepoint, where evo represents the evolved variant coverage and anc represents the ancestral variant coverage resulted on the vcf file. We also used directly the fastq files from population sequencing to estimate the evolved variant fitness over the ancestral as follows. For each variant in the consensus list we generated 12 search terms. The 12 terms were 20-base strings containing either the ancestral or the evolved allele, in the forward or reverse orientation and for 3 5-base sliding windows centered around the variant position. For all search terms aggregate counts were generated per allele and timepoint. In both cases, since we are mainly interested in heterozygous mutations, we assume that when anc > evo the population is a mix of individuals with the ancestral allele in homozygosity and heterozygosity with the evolved allele only. Mutations for which anc = evo or anc < evo in at least one timepoint were suggestive of including a fraction of individuals homozygous for the evolved allele. Nevertheless, they were still included in the analysis as far as they resulted in at least 3 timepoints for which anc > evo, since that could be an artifact because of low locus coverage. We used linear regression to model the natural logarithm of heterozygotes over the ancestor (2*evo/(anc-evo)) over time, where the slope represents the fitness coefficient of the variant in heterozygosity. Linearity was assessed with the Durbin-Watson statistic.

## FIGURE LEGENDS

**Figure S1. The barcode locus**. Barcoding was achieved via two consecutive transformations of a haploid version of our ancestor. The first transformation introduces a low complexity barcode, half of *URA3*, the KanMX marker and the gene for Cre recombinase under control of the Gal promoter, replacing the NatMX marker upstream of *GPA1*. The second transformation introduces a high complexity barcode and the rest of *URA3* gene via Cre induction.

**Figure S2. Mating efficiency depends on the haplotype**. Gene deletions (marked with KanMX) were introduced to the ancestor diploid (WT) and the resulting strains were subjected to sporulation and mass mating to a haploid with a NatMX marker. Mating efficiencies were estimated as the fraction of query diploids (colony-forming units (cfu) in double drug media) over the limiting mating partner marker (cfu on media with G418). Percentages of mating efficiencies of essentials over the non-essential *yur1Δ* of a representative experiment are shown. % recovery correlated with microcolony size observed upon tetrad dissection of the parental diploid. Correct marker integration and euploidy of the constructed strains were verified via tetrad dissection and observation of expected viability/marker segregation.

**Figure S3. Fitness of parental barcoded strains**. (A) Fitness of the parental strains (ancestor and evolved diploids) was assayed against a fluorescent version of the ancestor before and after low and high complexity barcode introduction. For most of the clones, fitness remained largely the same during barcoding. Killing activity phenotype is given for clones whose phenotype changed after transformation (see Table 1 for the phenotype of the rest). Double (low and high complexity) barcode derivatives were included in the barcode-based fitness assays. Asterisk indicates a clone that was omitted in the barcode-based assays. This clone had lost immunity during the last transformation. Ancestry of each clone can be traced with the annotations at the bottom. Color-coding reflects initial diploid. Each clone’s fitness was assayed four times. (B) Correlation between fluorescence- and barcode-based fitness estimates for each of the parental barcoded strains. Fluorescence-based fitness estimates are the same as in (A) for the low and high complexity barcode derivatives. Barcode-based fitness estimates are the same as in Figure 2.

**Figure S4. Fitness distributions of diploids diploid segregant pools from evolved parents F04-D and H06-C in the experimental and 3 alternative conditions**. The assays were barcode-based and the fitness was estimated with the algorithm published in (Venkataram et al. 2016). Fitness correlations are shown for 2 replicates per clone. Ancestor and evolved parents are annotated in cyan and orange, respectively. Derived diploids are annotated in grey. Pearson correlations for each replicate pair is shown at the top left of each panel.

## DATA AVAILABILITY

The raw short-read sequencing data reported in this paper are deposited under accession no. PRJNA775967 in the NCBI BioProject database.

## ACKNOWLEDGEMENTS

Ryan Vignogna for help with the design of CRISPR/Cas9 replacement constructs, and David Amberg and Brian Haarer for the providing hemizygotous gene deletion strains. Portions of this research were conducted on Lehigh University’s Research Computing infrastructure partially supported by NSF Award 2019035. This study was supported by grants from the National Institutes of Health R01GM127420 (G.I.L.) and R01AI164530 (S.F.L), and by the National Institute of Standards and Technology (S.F.L) and the Department of Energy (S.F.L).

