## Supplementary figures and images for "Overdominant and partially dominant mutations drive short-term adaptation in diploid yeast"

### Supplemental Figures

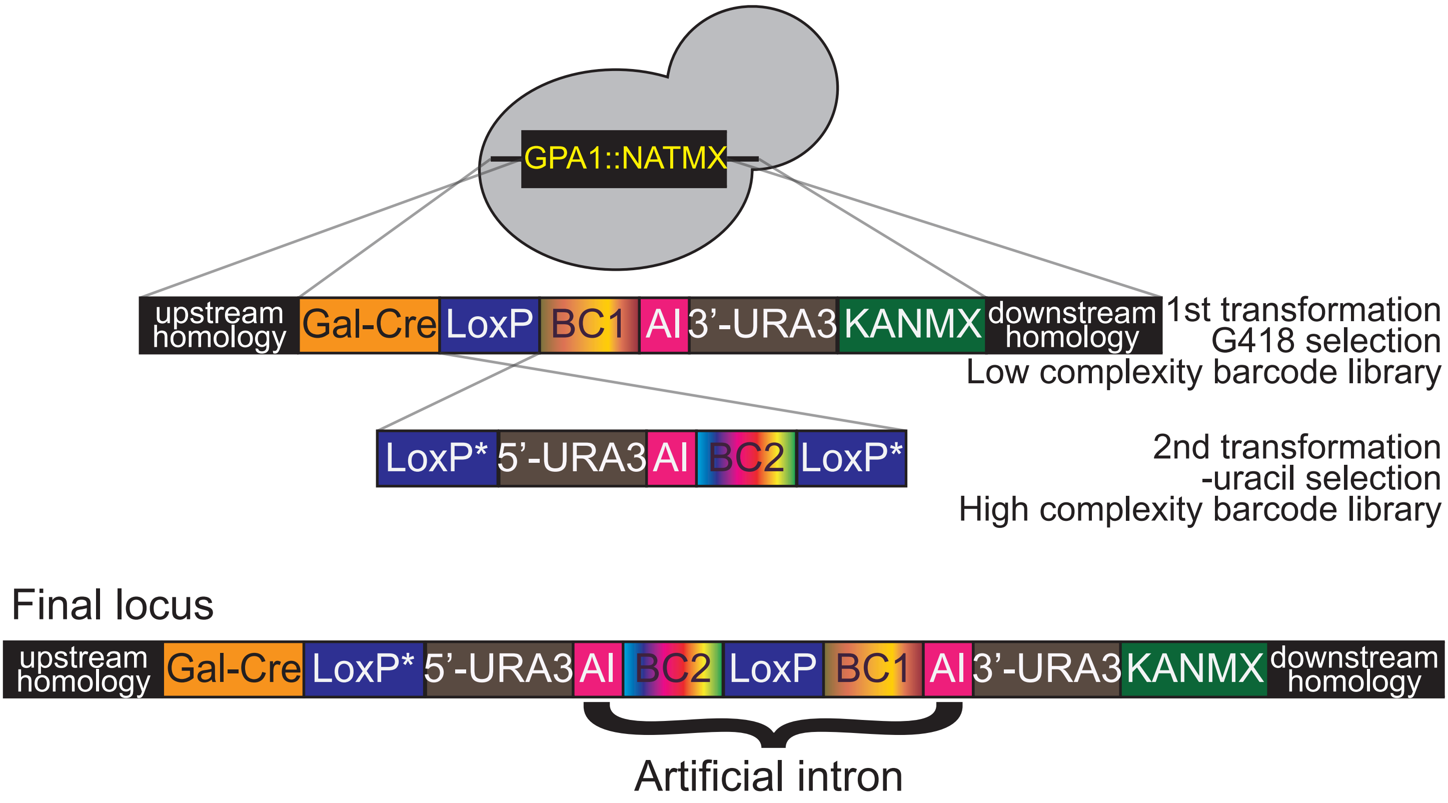

Figure S2

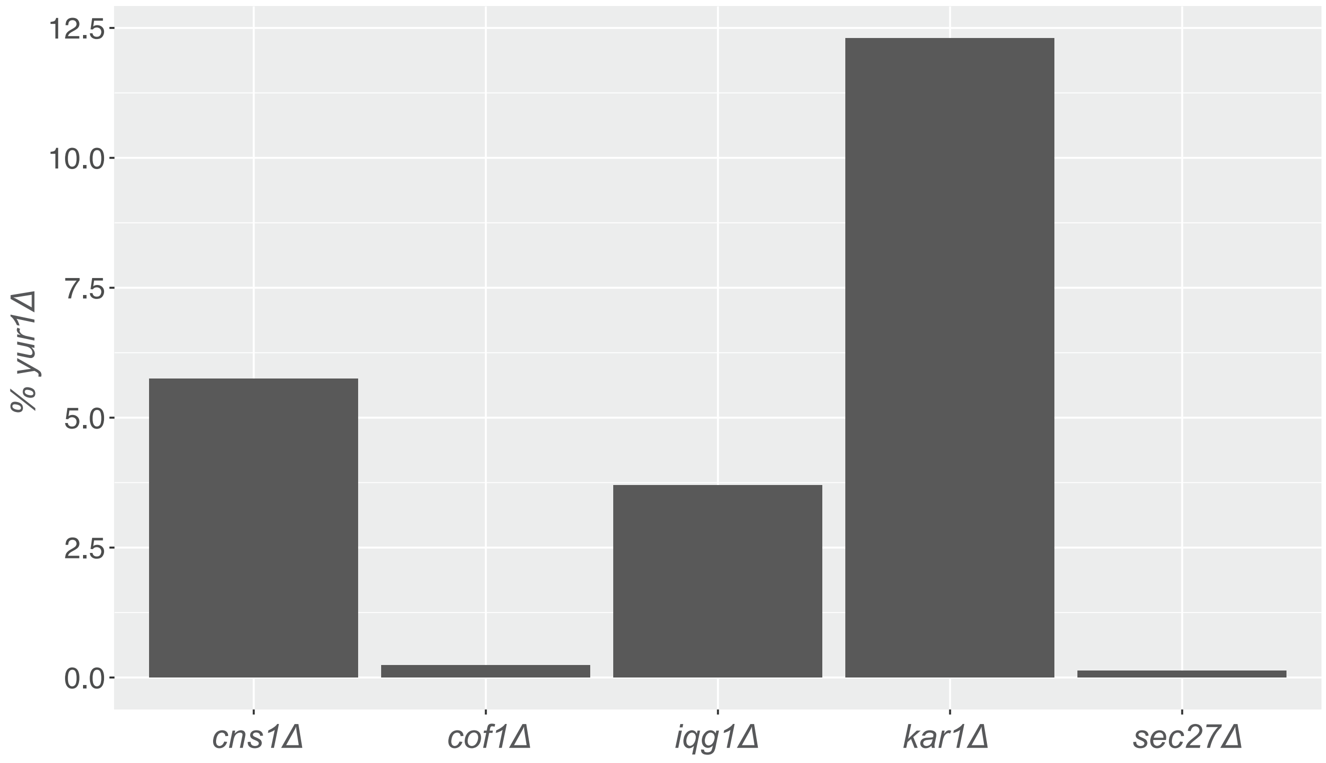

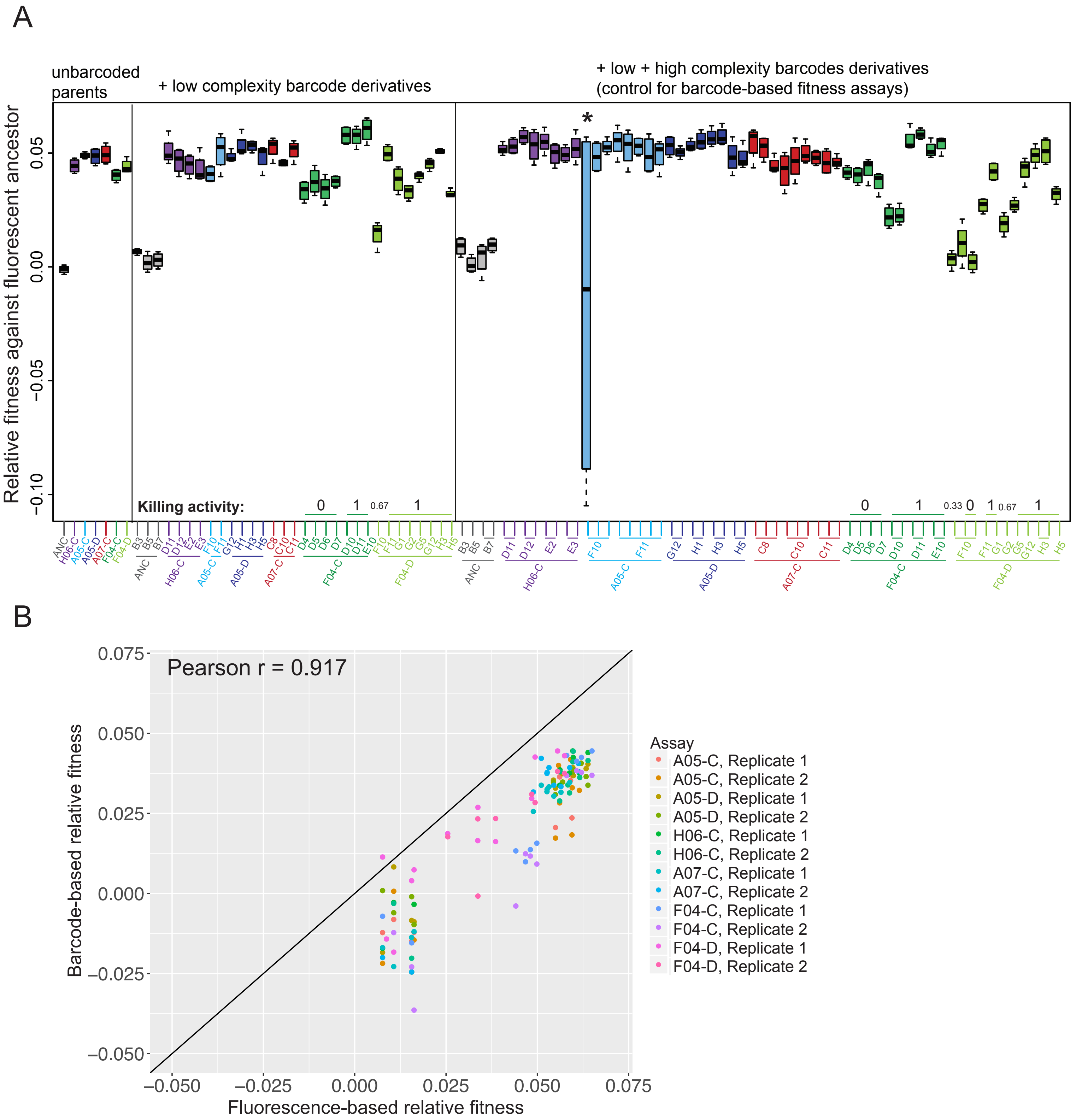

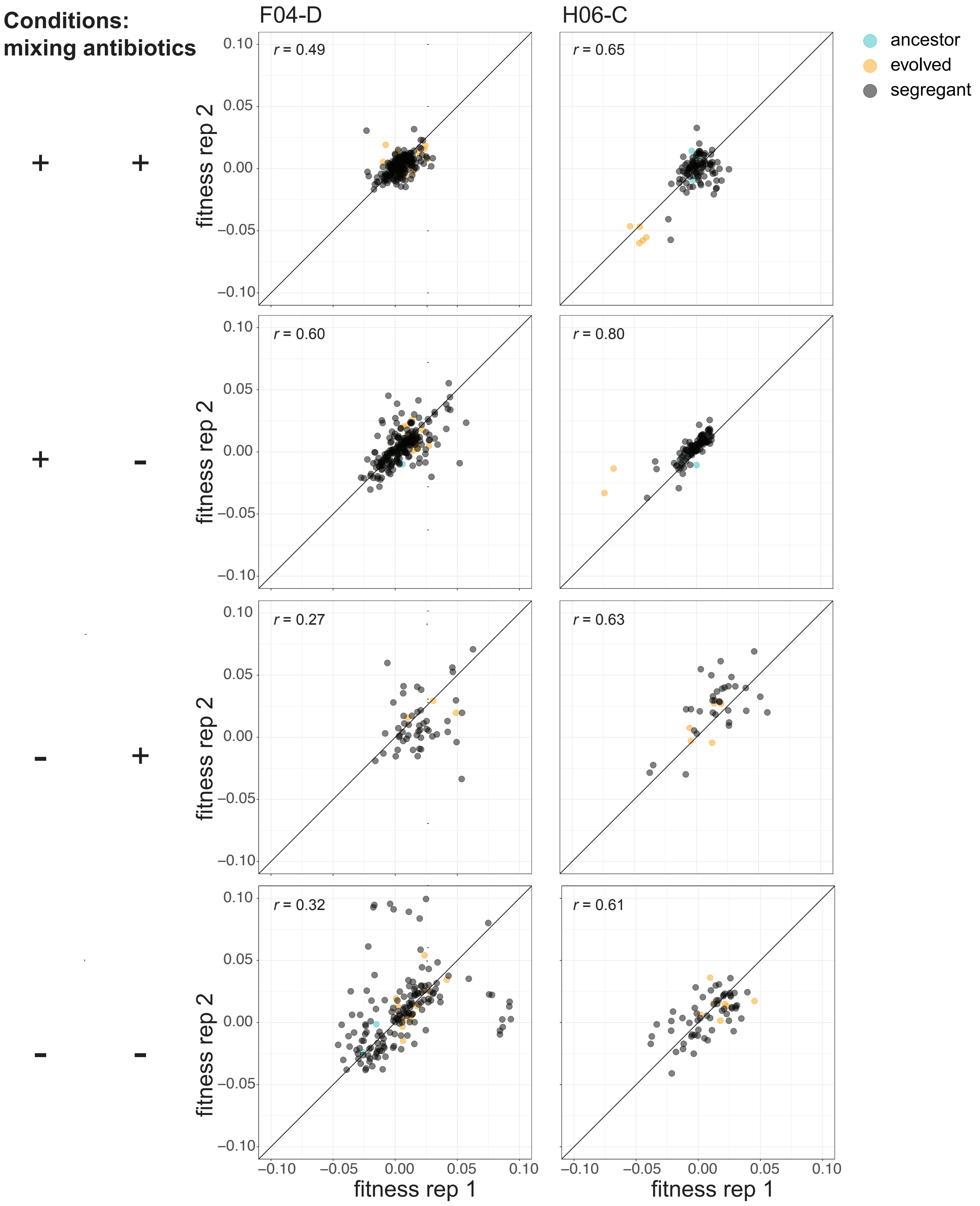
